# An efficient lentiviral CRISPRi approach to silence genes in primary human monocytes

**DOI:** 10.1101/2020.12.23.424242

**Authors:** Bingxu Liu, Matteo Gentili, David Lieb, Thomas Eisenhaure, Darrell J. Irvine, Nir Hacohen

## Abstract

Human primary monocytes are critical in controlling immune responses and tissue homeostasis. However, to identify and study molecular components that underlie the function of these cells, there remains a need to develop methods to perturb genes in these cells. Here we report lentiviral-based delivery of dCas9-KRAB and guide RNAs to efficiently silent target genes, such as *CD45* and *CD209*. We further show that sgRNAs against *TICAM1* dampen proinflammatory cytokine and interferon expression in response to lipopolysaccharide. When delivered by lentivirus, sgRNAs are incorporated into the genome, thus enabling pooled screening to perturb and identify coding and non-coding elements that contribute to the functions of primary human monocytes.

## Introduction

Monocytes play a central role in sensing pathogens and tissue damage and controlling the immune response and tissue homeostasis. The study of primary immune cells, including monocytes, has been feasible because of the availability of human peripheral blood. Monocytes and their differentiated progeny such as monocyte-derived dendritic cells have played important roles in studies of the association between genetic variants and the immune response,^1^ host-pathogen interactions,^2^ and identifying important pathogen restriction factors against pathogens like HIV.^3^ In addition to applications in basic immunology, monocyte-derived cells have been loaded with tumor antigens to create cell-based vaccines to activate neoantigen specific T cell responses against tumors.^4^ A key limitation in studies of monocytes is the difficulty in modifying their function through knockdown or knockout of endogenous genes.

While siRNA^5^ and shRNA^6^ libraries have been used in human monocytes, the high false-positive and false-negative rates continue to hinder large-scale application.^7,8^ In contrast, the CRISPR-Cas9 system enables efficient genetic perturbation including pooled screening for genome-wide analysis in a range of cell types, enabling new insights into diverse issues such as the role of genes in cancer cell biology^9^ or the role of cellular factors in host-pathogen interactions^10^. CRISPR can be used to reduce gene function either by cutting DNA with Cas9 or repressing gene expression with dCas9-KRAB.^7^

Despite advances in genetic perturbation methods and the growing demand for high-efficiency perturbation systems in monocytes, delivery of the CRISPR system into primary monocytes remains challenging. Here, we introduce a lentiviral method to allow efficient CRISPR interference (CRISPRi) for gene knockdown with minimal side effects in monocytes.

## Results

Post-mitotic monocytes are known to be resistant to lentivirus infection due to the anti-viral gene *SAMHD1*. Transduction of cells with a lentivirus together with a VLP containing the HIV-2-derived virion-associated VPX protein leads to degradation of SAMHD1 and efficient lentivirus integration.^11,12^ When we infected purified blood-derived CD14^+^ monocytes with VPX-carrying VLPs together with a lentivirus expressing KRAB-dCas9-P2A-mCherry (Fig 1a), we observed dCas9 levels increase in monocyte-derived DC (monocytes differentiated with GM-CSF and IL-4) over time and peak at day 6 with >70% mCherry+ (Fig 1b), consistent with previous reports.^11,12^

**Figure 1.**
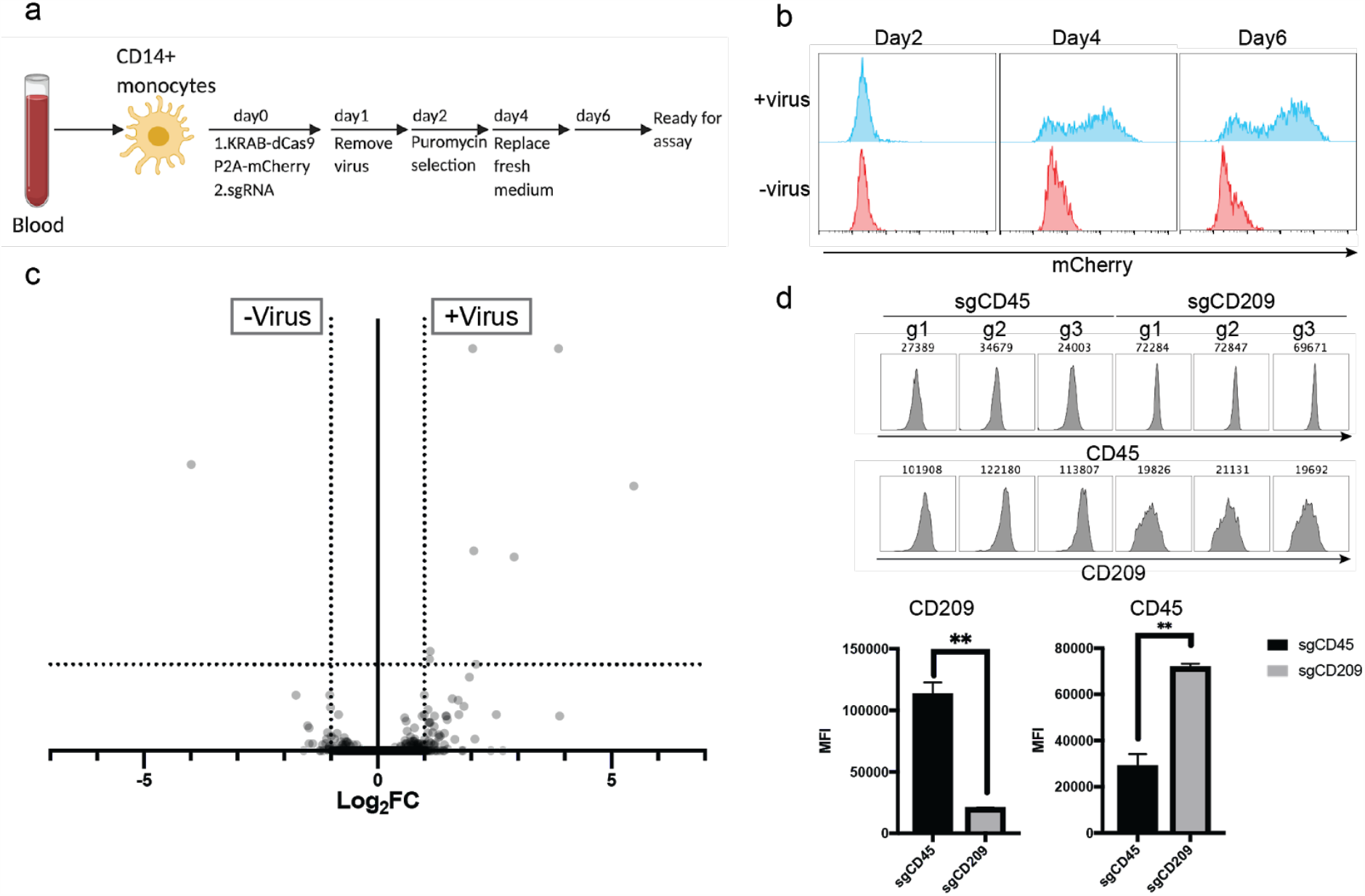
Lentiviral infection and CRISPRi-mediated knockdown of genes in MoDCs. a) Schematic of the transduction process. b) Lentiviral infection efficiency in MoDCs. Blue, cells transduced with virus; red, uninfected cells. c) Differential gene expression between transduced cells (+virus) and uninfected (-virus) cells stimulated with 50ng/ml LPS. X-axis shows fold change of gene expression; Y-axis shows significance of differential expression. Horizontal dotted line marks *p*=0.01. d) CD45 and CD209 expression of MoDCs transduced with 3 sgRNAs against *CD45* (sgCD45) or 3 sgRNAs against *CD209* (sgCD209). Histograms show distribution of fluorescence for bound antibodies targeting CD45 or CD209 protein, with the geometric mean value of the fluorescence intensity. Bar graphs show geometric MFI (mean fluorescence intensity) of cells transduced with sgRNAs against *CD45* or *CD209*, t test, **:p<0.005.

Lentivirus infection can trigger innate immune activation in monocytes. In order to understand if the lentivirus infection process influences cell states of monocyte-derived cells or their response to immune stimuli, we compared the transcriptome of the monocytes with or without lentivirus transduction in LPS-stimulated or resting conditions on day 7. Only a few genes showed significant differential expression between uninfected MoDCs and MoDCs transduced with dCas9 and sgRNA lentivirus in the resting state (Supplementary Figure 1) or following stimulation with LPS (Fig 1c and Supplementary Fig 2), suggesting delivery of the CRISPRi system through lentivirus did not alter the phenotype of monocyte-derived cells.

In order to quantify the efficiency of the CRISPRi system, MoDCs were infected with a lentivirus expressing sgRNAs against *CD45*, a constitutively expressed gene, or *CD209* which is induced during differentiation. On day 7, when cells were stained with anti-CD45 and anti-CD209 antibodies, MoDCs transduced with sg*CD45* exhibited a ∼70% reduction in CD45 while cells transduced with sg*CD209* had an ∼80% reduction in CD209, demonstrating the efficacy of the CRISPRi system in knocking down gene expression in these cells (Fig 1d).

Next we tested whether the CRISPRi system can be applied for the study of innate immune pathways. TLR signaling pathways sense a wide-range of pathogen-derived and damage-associated molecules, and are critical in the control of infections,^13^ autoimmunity^14^ and vaccine adjuvant design^15^. MYD88 and TRIF (encoded by the *TICAM1* gene) are two required downstream signal transducers of TLRs. While most TLRs, except TLR3, depend on MYD88 but not TRIF for proinflammatory cytokine production,^16^ TLR4 activation induces mixed transcriptional responses through MYD88 when it is on the plasma membrane and TRIF when it translocated to the endosome.^17^ *TICAM1* has been shown to be essential for interferon induction, while its contribution to production of the proinflammatory cytokine TNF-α has been controversial in different mouse cell models.^18-20^ Experiments using siRNAs against *TICAM1* have shown that TNF-α production in siTICAM1-transfected MoDC is dramatically lower than siCTRL transfected MoDCs.^5^

In order to test the consistency between CRISPRi and established siRNA methods, we transduced MoDC with sgsgTICAM1 or a non-targeting sgRNA (sgCTRL), RNA-seq analysis of resting MoDCs with sgTICAM1 or sgCTRL revealed that sgTICAM1 showed high specificity and efficiency since *TICAM1* was the only significantly downregulated gene (Fig 2a). Stimulation with TLR1/2 agonist Pam3CSK4 or TLR7/8 agonist R848 showed no difference in TNF-α production between the sgTICAM1 cells and the sgCTRL cells, indicating sgTICAM1 does not significantly influence inflammatory cytokine production through TLRs that depend on MYD88. However, MoDCs transduced with sgTICAM1 showed a dramatic reduction of TNF-α protein production compared to sgCTRL cells, indicating that TRIF is one of the important signal transducers of TLR4-mediated proinflammatory cytokine expression (Fig 2b). Further RNA-seq analysis revealed that the RNA levels of *TNF* together with other cytokine genes like *IFNB1* are also reduced in MoDCs transduced with sgTICAM1 (Fig 2c), suggesting that the TNF-α reduction effect at least partially come from TNF RNA level regulation. The results using the CRISPRi system in monocyte-derived cells showed high consistency with the established siRNA method.

**Figure 2.**
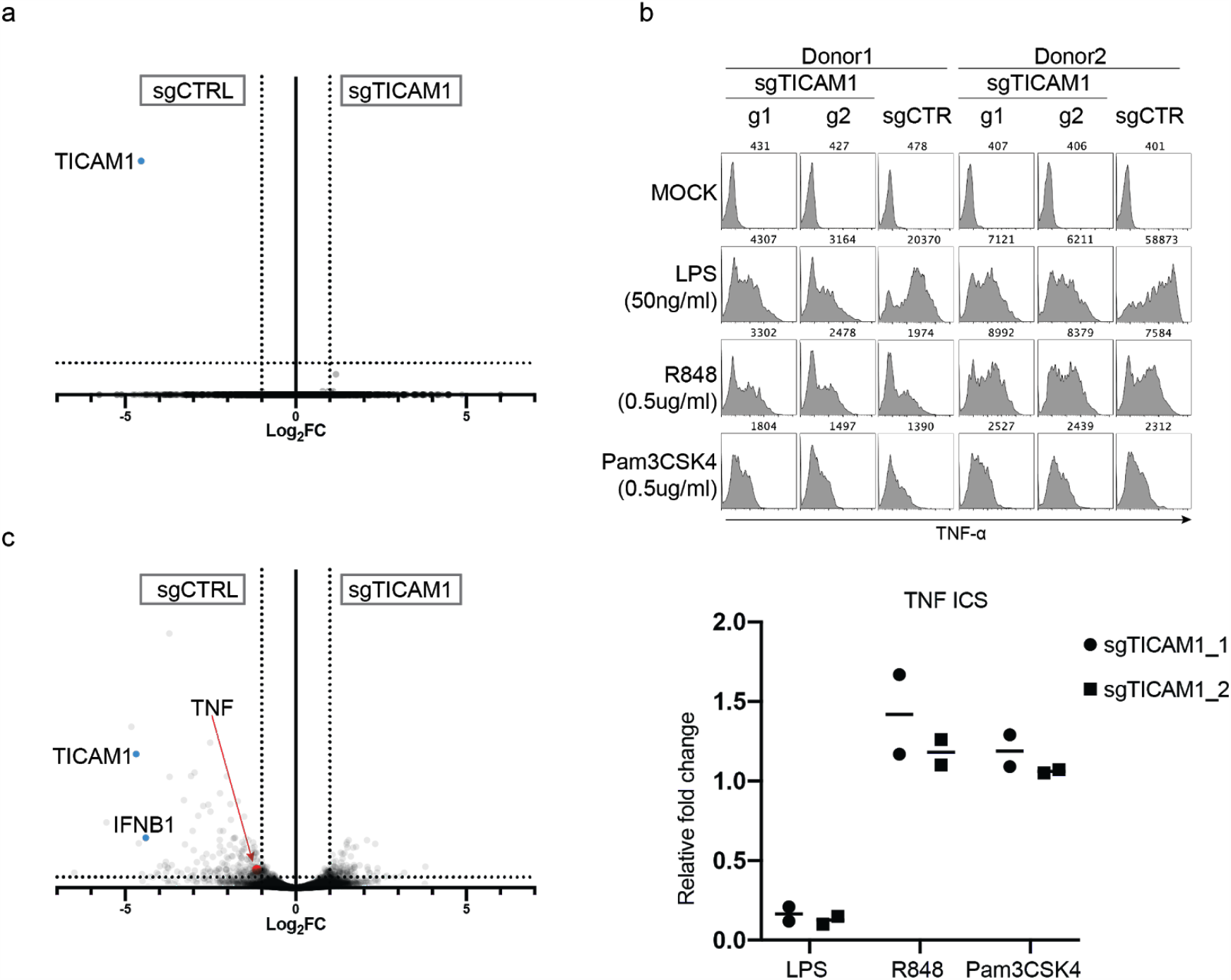
Changes in MoDCs transcriptomes in cells transduced with sgTICAM1 or sgCTRL. a) Transcriptome comparison of MoDCs transduced with sgTICAM1 guide 1 versus sgCTRL. b) Anti-TNFα intracellular staining of MoDCs transduced with sgTICAM1 and sgCTRL stimulated with medium(mock) or indicated stimuli for 5 hours. Top, histograms show distribution of fluorescence for bound antibodies TNFα, with the geometric mean value of the fluorescence intensity. Bottom, MFI normalized to sgCTRL. c) Transcriptome comparison of MoDCs transduced with sgTICAM1 guide 1 versus sgCTRL stimulated with 50ng/ml LPS for 4 hours. X-axis shows fold change of gene expression; Y-axis shows significance of differential expression. Horizontal dotted line marks p=0.01.

## Discussion

With >100 million CD14+ monocytes available for isolation from a single donor buffy coat, our method makes it possible to carry out genome-wide CRISPR screens (which typically require 40-50 million cells per replicate) to identify genes that affect almost any monocyte phenotype. In addition, combining CRISPR perturbation with single-cell RNA-seq is also now feasible, allowing reconstruction of genetic networks as was done in mouse bone-marrow-derived dendritic cells upon LPS stimulation.^21^ Another important problem is to understand the role of non-coding elements in the human immune response.^1^ While eQTL provides associations, it does not demonstrate causality, which CRISPRi can do by mapping enhancer-promoter connections.^22,23^ Hence, this method will enable systematic functional analysis of causal variants that contribute to immune response variation in the human population.

## Method

Virus packaging.0.8 million 293T cells were seeded in each well of a 6 well plate with 1ml of DMEM. For dCas9 and sgRNA encoding virus, cells in each well are transfected with 0.4ug pCMV-VSV-G, 1ug PsPAX2 and 1.6ug of pHR-SFFV-KRAB-dCas9-P2A-mCherry(addgene, #60954) or agOpti (addgene,#85681) using TransIT-293(MirusBio). On day 2, replace fresh RPMI. Supernatant is harvested 30 hours after media replacement.

### Lentivirus infection

CD14+ monocytes were isolated from adult human blood (Research Blood components)^2^. 100,000 Monocytes in 50 ul MDDC medium (RPMI with Glutamax, 10% FBS (Gibco), PenStrep(Gibco), 50ug/ml Gentamicin(Gibco) and 0.01M HEPES(Gibco)) were infected with 100 ul KRAB-dCas9 virus, 50 ul sgOpti virus and 50ul Vpx VLP in the presence of 10ng/ml recombinant human GM-CSF(Miltenyi) and 50ng/ml IL-4 (Miltenyl) and 8ug/ml Protamin(Sigma). On day 1, the monocytes were pelleted and resuspended in fresh MDDC medium with GM-CSF and IL-4. On Day 2, 2ug/ml puromycin were applied for selection. On Day4, the cells were pelleted and resuspended in the fresh MDDC medium with GM-CSF and IL-4. The cells were assayed on Day 7. For Monocyte-derived Macrophage, 25ug/ml M-CSF is used to replace GM-CSF and IL-4.

### Staining

For cell surface staining, cells were collected and washed and stained in FACS buffer (PBS, 1% BSA, 1mM EDTA, 0.01 NaN3). Human DC-SIGN/CD209 PE-conjugated Antibody (R&D, #FAB161P-025) and anti-human CD45 AF700 (Biolegend, #304024) are for MDDC surface staining. For intracellular cytokine staining, cells were incubated with GolgiPlug 30min before adding TLR agonists. After 5 hours TLR ligands stimulation, the cells were fixed and permeabilized using BD Fixation and Permeabilization Solution Kit with BD GolgiPlug (Cat. No. 555028) and stained with APC anti-human TNF-α Antibody (BioLegend, #502913).

### RNA-seq

The cells were cultured with or without LPS for 4 hours, Cells were pelleted and resuspended in ice-cold MACS buffer(PBS, 1% BSA, 1mM EDTA), 200-500 mCherry positive cells were sorted in 20 ul Lysis buffer per well of a 96 well plate, and cDNA libraries produced using a modified Smartseq2 protocol^24^. DEseq2 was used to find differentially expressed genes.

## Author contribution

B.L, M.G, D.J.I and N.H planned and designed experiments. B.L, M.G, D.L, and T.E performed experiments and analyzed data. B.L, D.J.I and N.H wrote the manuscript.

