## Supplemental materials for "An efficient lentiviral CRISPRi approach to silence genes in primary human monocytes"

**Supplement Figures**

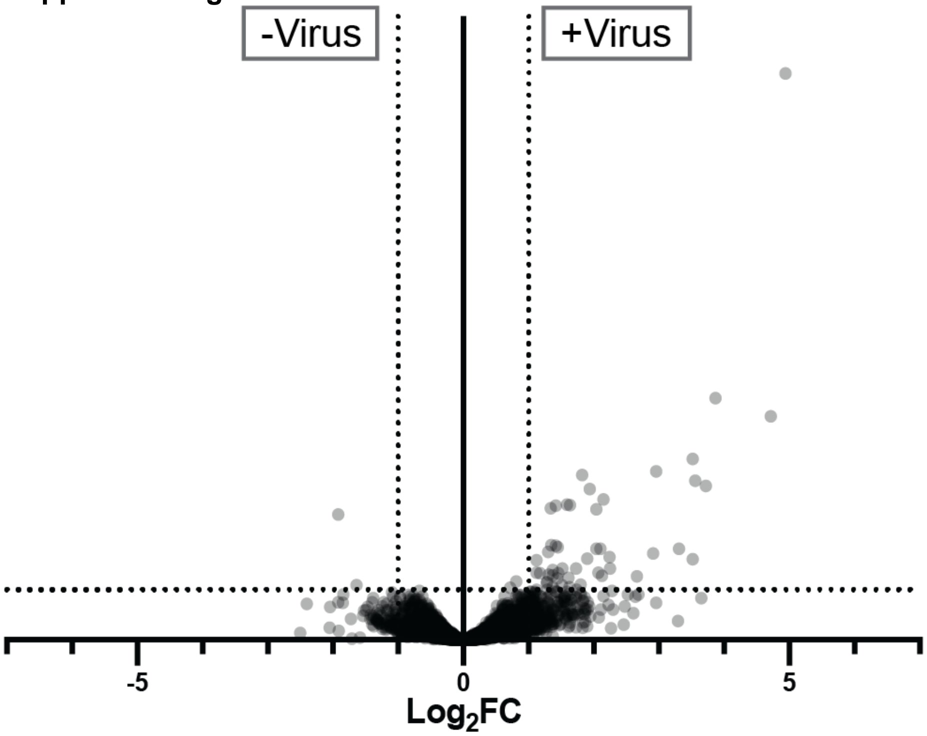


**Supplementary Figure 1**. Differential gene expression between transduced MoDCs (+virus) and uninfected (-virus) cells. X-axis shows fold change of gene expression; Y-axis shows significance of differential expression. Horizontal dotted line marks $p$=0.01.

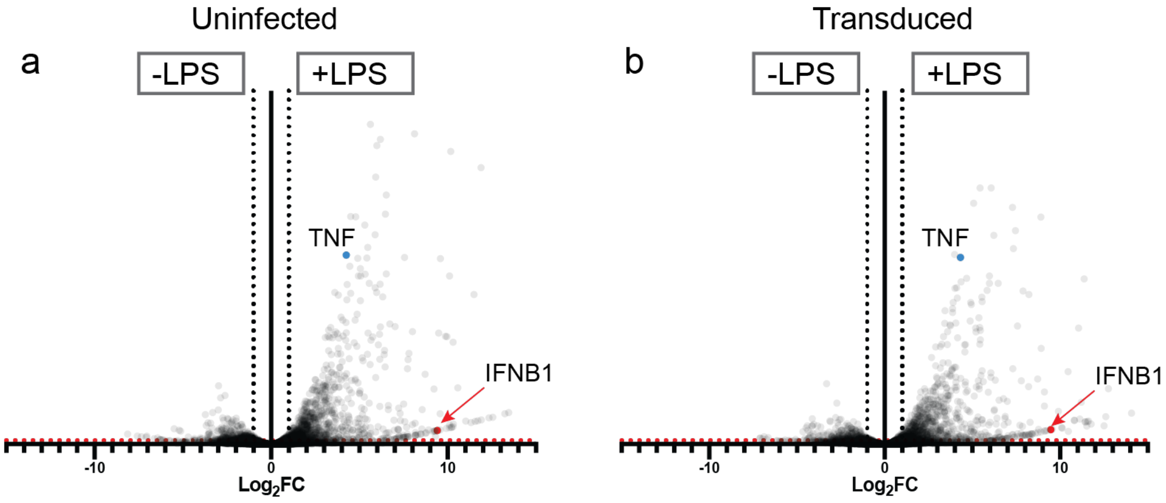


**Supplementary Figure 2**. a) Differential gene expression between unstimulated and 50ng/ml LPS stimulated cells stimulated uninfected MoDCs. b) Differential gene expression between unstimulated and 50ng/ml LPS stimulated transduced MoDCs. X-axis shows fold change of gene expression; Y-axis shows significance of differential expression. Horizontal red dotted line marks $p$=0.01.
